# Proteomic characterization of extracellular vesicles from 12 commensal bacterial species

**DOI:** 10.1101/2025.11.06.686325

**Authors:** Roselydiah Nasipwondi Makunja, Arina Maltseva, Tiina Pessa-Morikawa, Masuma Khatun, Maria Stensland, Mari Heinonen, Anna Kaisanlahti, Subhashini Muhandiram, Justus Reunanen, Terhi Ruuska-Loewald, Tuula A. Nyman, Mikael Niku

## Abstract

Bacterial extracellular vesicles (bEVs) produced by intestinal commensal bacteria mediate host-microbe interactions, but their proteomes have been explored in only a small number of species. We characterized bEVs from *in vitro* cultures of 12 species of Actinomycetota, Bacillota (Firmicutes), Bacteroidota, Fusobacteria, Pseudomonadota, and Verrucomicrobia. This is the first report of bEVs from *Veillonella magna*, an exceptional gram-negative Bacillota, and the gram-positives *Peptostreptococcus russellii* and *Turicibacter sanguinis*. The morphology and protein subcellular localization patterns of bEVs reflected the envelope structures of their parent bacteria. Notably, *V. magna* may be able to produce both outer membrane and cytoplasmic membrane vesicles. The proteome compositions were dictated by phylogeny, suggesting largely non-selective packaging of proteins. Annotation of protein functions indicated roles in nutrient metabolism and transport, and in host immune system modulation. Peptidoglycan modifying enzymes and the abundant bacteriophage proteins in *Enterobacter cloacae* and *Limosilactobacillus reuteri* bEVs may be involved in vesicle biogenesis. The functions of many abundant bEV proteins are unknown. The most abundant bEV proteins were largely species-specific, but we identified several conserved proteins that may be used as markers to distinguish commensal bEVs from host EVs. Comparison to previous *in vitro* and fecal metaproteomics data indicates that bEV proteome compositions are reproducible.

## Introduction

Bacterial extracellular vesicles (bEVs) are membrane-derived lipid bilayer nanoparticles, secreted by both gram-negative and gram-positive bacteria, ranging in size from 40 to 400 nm (1). Their biogenesis is influenced by the structure of the cell envelope, which differs between gram-negative and gram-positive bacteria (2, 3). In gram-negative bacteria, bEVs are secreted by outer membrane blebbing or phage-mediated explosive cell lysis, while in gram-positive bacteria, they are produced through budding or blebbing of the cytoplasmic membrane or phage-mediated bubbling cell death (1). The bEV cargo consists of proteins, nucleic acids, lipids, and small-molecular metabolites, which may be delivered to other microbial or host cells (4, 5). Compared to simple diffusion, bEVs can deliver highly concentrated signals over extended distances, referred to as quantal secretion (1). During transit, the cargo is protected from environmental inactivators and degrading enzymes (6).

In the intestine, commensal bEVs play diverse roles in bacteria-bacteria and bacteria-host interactions (7–9). They aid bacteria in nutrient acquisition, niche competition, and adaptation to environmental stress (10). bEVs carry enzymes that degrade complex carbohydrates, supporting the growth and survival of both the producer and other members of the microbial community (11). bEVs prime the host immune system against viral (12) and bacterial infections (13); they may have anti-inflammatory properties (14–16) and improve epithelial barrier integrity (14, 17). The immunomodulatory effects of bEVs may be even more potent than those of the producing bacteria since they can be internalized by host cells, thereby activating intracellular immune receptors (12, 18). Thus, bEVs hold promise for therapeutic applications, including immunotherapy, management of inflammatory bowel disease (IBD), cancer treatment, and drug delivery (4, 19, 20).

Commensal bEVs and especially their proteomes have only been characterized from a small part of the intestinal microbiota, with most studies focusing on a few bacterial species such as *Akkermansia muciniphila* (21–26), *Bacteroides fragilis* (18, 27–29), and *Bifidobacterium longum* (30, 31). They are usually studied *in vitro*, as *in vivo* research is complicated by a lack of markers for distinguishing bacterial from host extracellular vesicles (EVs) (4, 32). The growth conditions, bacterial strains, and bEV processing workflows may significantly influence the observed bEV composition (4, 19, 32, 33).

In the current study, we assessed the structures and proteomes of bEVs derived from gram-negative and gram-positive commensal bacteria, representing all major phyla and orders of the mammalian intestinal microbiota. We searched for conserved proteins that could serve as markers for bEVs in animal samples. To assess the effects of bEV processing workflows and experimental methods on the observed bEV characteristics, we compared our data to previous bEV studies from the same bacterial species.

## Materials and methods

The experimental protocol has been submitted to the EV-TRACK knowledgebase (EV-TRACK ID: EV250029) (34).

### Bacterial culture

The bacterial strains (Table 1) were obtained from culture collections, except for *Lactobacillus amylovorus* and *Limosilactobacillus reuteri*, which were in-house isolates from sow feces collected from Finnish pig farms (35). The cultured strains were verified by MALDI-TOF and/or Sanger sequencing of the 16S rRNA genes.

**Table 1.**
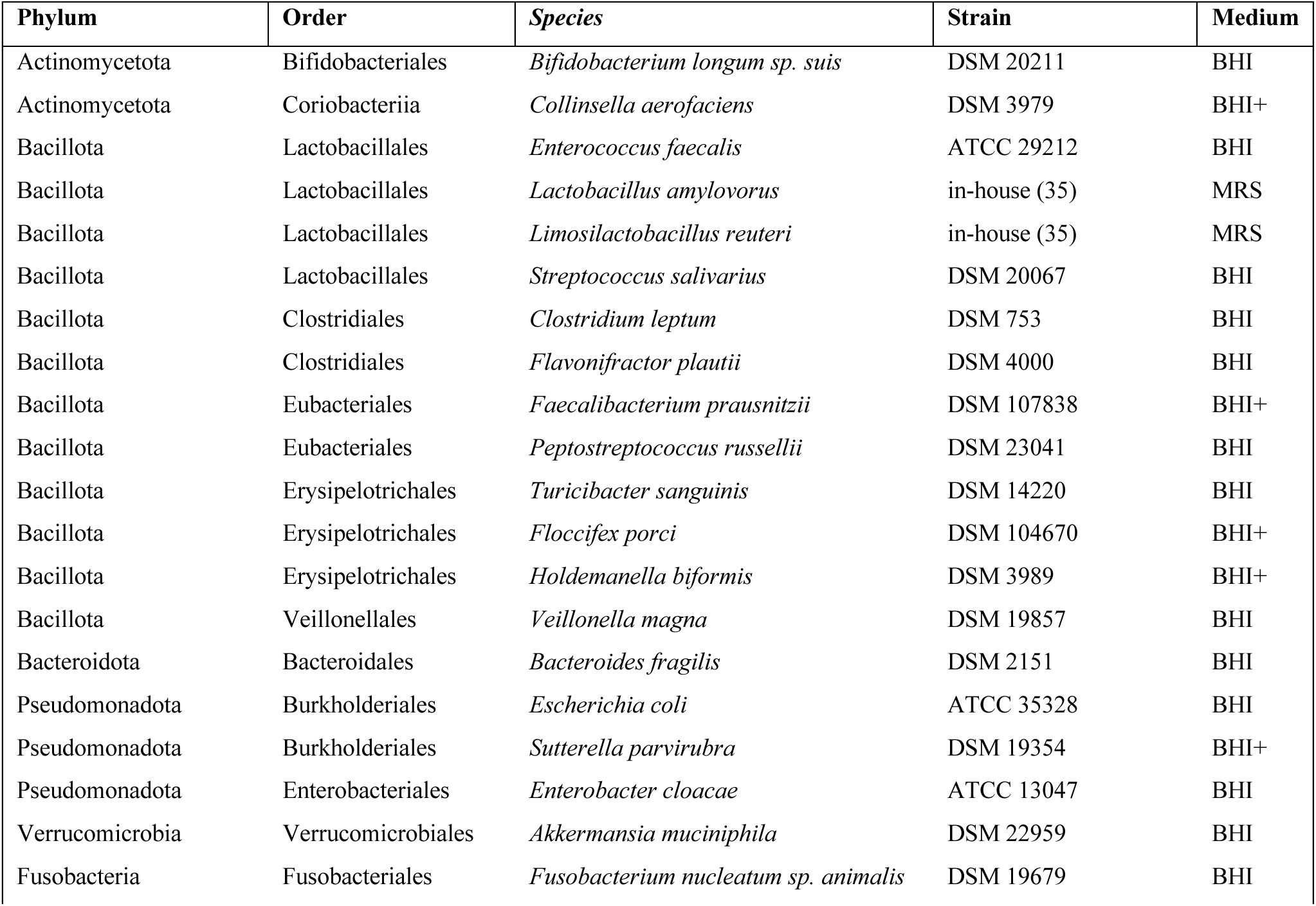
Bacterial strains and culture media. BHI: Brain Heart Infusion broth (BHI). BHI+: BHI with supplements. MRS: deMan, Rogosa and Sharpe broth. The Bacillota phylum was previously Firmicutes.

The bacteria were cultured anaerobically at +37 °C, without shaking (Table 1). Most of the bacteria were cultured in the Brain Heart Infusion broth (BHI; Merck Millipore #53286). For some species indicated in the BHI medium was supplemented with yeast extract (5 g/l), hemin (5 mg/l), vitamin K (1 mg/l), cellobiose (1 g/l), maltose (1 g/l), and cysteine (1 g/l). *L. reuteri* and *L*. *amylovorus* were cultured in deMan, Rogosa and Sharpe broth (MRS; Oxoid / Thermo Fisher Scientific #CM0359). We also tested Gifu Anaerobic Medium (GAM; Nissui Pharmaceutical #05433), but it contained high concentrations of EV-like particles and filtration resulted in poor bacterial growth.

For the final 30 ml cultures (each species with three replicates) used for EV isolation, the culture media were pre-filtered using 0.45 µm and 0.1 µm sterilization filters (Acrodisk Supor, Pall Lab/Cytiva) to deplete EV-like particles. The EVs were collected when the cultures were clearly cloudy (depending on bacterial strain, 1–4 days).

The following strains did not grow in our experimental conditions: *Alistipes senegalensis* (DSM 25460), *Anaerostipes hadrus* (DSM 3319), *Eubacterium biforme* (DSM 3989), and *Prevotella histicola* (DSM 26979).

### Isolation of bacterial extracellular vesicles

Bacterial cells were removed by centrifugation for 10 min at 3100 × *g* (4000 rpm) using an Eppendorf 5810 R Centrifuge, with swing-bucket rotor A-4-81. The supernatants were filtered through a 0.22 µm sterilization filter (Millex-GP, Merck Millipore) using a peristaltic pump and stored overnight at +4 °C. EVs were collected by ultracentrifugation of 3 × 22 ml culture supernatant per species at 110000 × *g* for 2 hours at +4 °C, using Optima LE-80K ultracentrifuge with a Ti 50.2 rotor, k-factor 143.3 (Beckman Coulter), and resuspended in 500 µl phosphate-buffered saline (PBS, 0.22 µm pre-filtered). Size exclusion chromatography was performed using 70 nm qEV Original columns (Izon Science). 500 µl per sample was loaded, and 3 × 0.5 ml fractions were collected by washing with PBS. Finally, samples were concentrated by ultrafiltration at 3200 × *g* using 10 kDa MWCO 4 ml Amicon Ultra tubes (Merck), loading 1.5 ml per sample and collecting 100 µl.

### Characterization of bacterial extracellular vesicles by NTA and electron microscopy

Particle concentration and size were measured with the ZetaView PMX-120 nanoparticle tracking analysis (NTA) instrument (Particle Metrix GmbH, Ammersee, Germany) equipped with a Z NTA cell assembly, a blue (488 nm, 40 mW) laser, and a CMOS camera with 640 x 480-pixel resolution. Samples were diluted in a total volume of 1 ml of particle-free ultra-pure milli-Q water to obtain 50–200 particles per frame. Videos in NTA mode were recorded at 11 positions across the measurement chamber in two-second increments at 30 FPS framerate with camera shutter speed at 100 s^-1^ and sensitivity at 85. The temperature was controlled at 22°C for NTA. Videos were processed, and outliers (>10% CV) were removed using the Grubbs method with the built-in ZetaView software (version 8.05.12 SP2). Particles between 10-1000 nm in diameter with a minimum trace length of 15 frames and a minimum brightness of 20 were included in the analysis. Culture media without bacteria were used as negative controls (nine replicates of BHI medium, out of which one replicate was removed as an outlier due to much higher particle concentration and different size range; and three replicates of MRS medium).

EVs were prepared for electron microscopy (EM) as in Puhka *et al*. (36) by loading to carbon-coated and glow-discharged 200 mesh copper grids with a pioloform support membrane. EVs were fixed with 2% paraformaldehyde in NaPO4 buffer, stained with 2% neutral uranyl acetate, and further stained and embedded in uranyl acetate and methyl cellulose mixture (1.8/0.4 %). EVs were viewed with transmission EM (TEM) using Jeol JEM-1400 (Jeol Ltd., Tokyo, Japan) operating at 80 kV. Images were taken with Gatan Orius SC 1000B CCD-camera (Gatan Inc., USA) with 4008 × 2672 px image size and no binning.

### LC-MS/MS analysis of bacterial extracellular vesicle proteomes

For the isolation and preparation of bEV proteins, a magnetic bead-based method was used for most species. Here, the EV proteins were precipitated with 70% acetonitrile onto magnetic beads (MagReSynAmine, Resyn Biosciences), and then washed on the beads with 100% acetonitrile, 70% ethanol, and then resuspended in 100 µl of 50 mM ammoniumbicarbonate. The proteins were reduced by the addition of 0.5M DTT to a final concentration of 10 mM and incubation at 56°C for 30 min. To alkylate proteins, 2.7µl of 550 mM iodoacetamide (IAA) was added to a final concentration of 15 mM, and samples were incubated at room temperature in the dark for 30 min. 0.5 µg trypsin was added to each sample for overnight on-beads protein digestion at 37°C. The resulting peptides were concentrated using the STAGE-TIP method with a C18 resin disk (Affinisep).

An earlier non-bead-based protocol was used for EV proteins of *E. faecalis, E. cloacae, L. amylovorus, L. reuteri, B. longum* sp. *suis* and *B. fragilis*. Here, 80 µl of each EV sample was mixed with ProteaseMax surfactant (Promega) in 50 mM ammonium bicarbonate to a final concentration of 0.1%. Samples were vortexed for one minute and heated at 95°C for five minutes. Protein reduction and alkylation were performed as reported above. Trypsin digestion was performed in solution.

The liquid chromatography and tandem mass spectrometry (LC-MS/MS) analysis was performed using a nanoElute UHPLC coupled to a timsTOFfleX mass spectrometer (Bruker Daltonics, Bremen, Germany) via a CaptiveSpray ion source. Peptides were separated on a 25 cm reversed-phase C18 column (1.6 µm bead size, 120 Å pore size, 75 µm inner diameter, Ion Optics) with a flow rate of 0.3 µl/min and a solvent gradient from 0-35% B in 60 min. Solvent B was 100% acetonitrile in 0.1% formic acid, and solvent A 0.1% formic acid in water. The mass spectrometer was operated in data-dependent Parallel Accumulation-Serial Fragmentation (PASEF) mode. Mass spectra for MS and MS/MS scans were recorded between m/z 100 and 1700. Ion mobility resolution was set to 0.85–1.35 V·s/cm over a ramp time of 100 ms. Data-dependent acquisition was performed using 10 PASEF MS/MS scans per cycle with a near 100% duty cycle. A polygon filter was applied in the m/z and ion mobility space to exclude low m/z, singly charged ions from PASEF precursor selection. An active exclusion time of 0.4 min was applied to precursors that reached 20,000 intensity units. Collisional energy was ramped stepwise as a function of ion mobility.

Protein identification was performed using the MaxQuant software (versions 2.1.3.0,2.4.3.0 and 2.4.7.0) (37). Parameters were set as follows: fixed modification: carbamidomethylation (C), protein N-acetylation, and methionine oxidation as variable modifications. The first search error window of 20 ppm, and mains search error of 4.5 ppm. Trypsin without proline restriction enzyme option was used, with two allowed miscleavages. Minimal unique peptides were set to 1, and the false discovery rate (FDR) allowed was 0.01 (1%) for peptide and protein identification. Species-specific databases were generated from proteomes downloaded from NCBI Genomes. In addition, human, bovine, and pig databases downloaded from Uniprot were included in the MaxQuant searches. The generation of reversed sequences was selected to assign FDR rates.

Most of the bacteria were analyzed in three culture replicates. For *F. nucleatum* sp. *animals*, *H. biformis, P. russellii*, *S. salivarius* and *T. sanguinis*, the replicates were pooled for LC-MS/MS analysis, due to low signal intensities.

### Prediction of protein cellular localization and function

The cellular localization of bEV proteins was predicted using PSORTb version 3.0.3 web server (38) and DeepLocPro (39). Analysis was performed for each species separately, using the default parameters and the species gram type. DeepLocPro predictions were used in cases where PSORTb predictions returned an “unknown”. Some of the 50S ribosomal proteins were erroneously predicted as outer membrane or extracellular by DeepLocPro; these were reclassified as cytoplasmic. Eight *V. magna* proteins were predicted by DeepLocPro as cell wall proteins; since gram-negative bacteria do not have an actual cell wall, and these proteins are known to span several envelope layers, they were reclassified as unknown.

Protein functions were predicted as clusters of orthologous groups (COG) using eggNOG-mapper v2 (40). Proteins without eggnog annotations were predicted using PANNZER2 (41), converting the Gene Ontology biological functions to COGs using a Python script by Szczerbiak *et al*. (42). Remaining unannotated proteins were processed using the NCBI CD-Search Tool (43).

### Comparative analysis of bacterial extracellular vesicle proteomes

To compare bEV proteome compositions across species, we defined protein orthogroups using OrthoFinder v2.5.5 (44) with default parameters. *F. nucleatum* was excluded from this analysis due to the small number of identified proteins. For the quantitative analyses, the remaining data were quantile normalized with the limma R package (45). For inference of the proteome similarity tree and the proteomic versus consensus phylogenetic trees comparison, orthogroup abundances were averaged across replicates by species, and orthogroups with averaged relative abundance values not reaching 0.1% in at least one species were removed to minimize the effect of the total number of identified proteins on tree topology. For the nonmetric multidimensional scaling (nMDS) ordination, orthogroups detected in less than 2 samples were excluded. Orthogroups detected in five out of seven gram-positive species or all of the four remaining gram-negative species were regarded as conserved. The lower threshold value in gram-positive species was chosen due to protein-poor bEV proteomes obtained for *B. suis*, *S. salivarius,* and *T. sanguinis* (less than 50 orthogroups identified). The Venn diagrams were created with the eulerr R package (46).

For the proteomic similarity tree, bEV proteomes averaged by species were clustered using the Jaccard distance matrix (vegdist function of the vegan package, method = “jaccard”, binary = FALSE), with complete linkage as the grouping algorithm (47) and plotted with the dendextend package (48). Branch support was assessed by approximately unbiased (AU) p-values using multiscale bootstrap resampling (49) with 5,000 iterations in the pvclust package (50). This number of iterations ensured accurate estimation of AU p-values (standard errors were less than 0.025). The phylogenetic tree for the tanglegram calculation was taken as the phy file from the NCBI Taxonomy Common Tree tool. Before comparison, phylogenetic and proteomic trees were made ultrametric using non-negative least squares with the phangorn package (51). The topological similarity between the proteomic and phylogenetic trees was evaluated using the Fowlkes-Mallows index (52). The index measures the similarity between two dendrograms. It quantifies how similarly the trees are partitioned into clusters, with values of the index ranging from 0 (no similarity) to 1 (perfect agreement). The index estimates the similarity of the compositions of clusters obtained when the trees are split at various numbers of clusters (k). The number of clusters (k) varies from 2 up to (N-1), where (N) is the total number of end branches in each tree (proteomic and phylogenetic). This allows comparison of the overall clustering structure between the two trees. Correspondence between inter-species distances on the trees was measured using the coefficient of cophenetic correlation (53). Fowlkes-Mallows index, cophenetic correlation value, and tanglegrams were calculated and plotted with the dendextend package (48).

The proteome ordinations were visualized using nMDS based on Euclidean distances among the samples (vegan package) (47). All plots were drawn with the ggplot2 package (54). Heatmaps were created with the ComplexHeatmap package (55). The values of molecular weight for detected proteins in proteomes were predicted with the Peptides package (56) to test possible correlations with bEV size or gram type.

### Comparison to fecal extracellular vesicle metaproteomic datasets

Peptide-level fecal EV proteome datasets (with protein identifications based on trEMBL) were obtained from the PRIDE repository (accessions PXD045755 and PXD047510) (57, 58). Peptides designated as human were removed. The remaining peptides were analyzed using UniPept (59). We included only those peptides that could be assigned to the genera listed in Table 2 (that is, assigned to the correct bacterial class and not assigned to another order, family, or genus). Average relative intensities were then calculated for unique protein groups. Samples with less than 10 identified protein groups were excluded from further analysis. Protein localizations were predicted from the amino acid sequences available in Uniprot or Uniparc, as described above.

**Table 2.** Bacterial extracellular vesicle diameters (nm; mean ± STDEV) and concentrations (particles/ml; mean ± STDEV). NTA: nanoparticle tracking analysis (NTA); TEM: transmission electron microscopy. Brain Heart Infusion broth (BHI) and deMan, Rogosa and Sharpe (MRS) culture media were used as negative controls. The Bacillota phylum was previously Firmicutes.

| Phylum | Species | Gram | NTA diam. | TEM diam. | NTA concentration |
| --- | --- | --- | --- | --- | --- |
| Actinomycetota | *Bifidobacterium longum sp. suis* | + | 131 ± 47 | 99 ± 56 | 1.6 × 10<sup>10</sup> ± 2.5 × 10<sup>9</sup> |
| Bacillota | *Enterococcus faecalis* | + | 146 ± 61 | 96 ± 34 | 1.1 × 10<sup>11</sup> ± 4.3 × 10<sup>10</sup> |
| Bacillota | *Limosilactobacillus reuteri* | + | 143 ± 55 | 110 ± 41 | 2.6 × 10<sup>10</sup> ± 8.5 × 10<sup>9</sup> |
| Bacillota | *Streptococcus salivarius* | + | 124 ± 53 | 65 ± 19 | 1.9 × 10<sup>9</sup> ± 5.1 × 10<sup>8</sup> |
| Bacillota | *Faecalibacterium prausnitzii* | + | 247 ± 105 | 70 ± 21 | 1.7 × 10<sup>12</sup> ± 2.9 × 10<sup>11</sup> |
| Bacillota | *Peptostreptococcus russellii* | + | 104 ± 39 | 73 ± 16 | 2.3 × 10<sup>10</sup> ± 1.9 × 10<sup>10</sup> |
| Bacillota | *Turicibacter sanguinis* | + | 134 ± 55 | 50 ± 13 | 6.5 × 10<sup>9</sup> ± 5.4 × 10<sup>9</sup> |
| Bacillota | *Veillonella magna* | – | 167 ± 66 | 74 ± 31 | 4.2 × 10<sup>11</sup> ± 1.9 × 10<sup>11</sup> |
| Bacteroidota | *Bacteroides fragilis* | – | 150 ± 63 | 56 ± 19 | 7.1 × 10<sup>9</sup> ± 2.1 × 10<sup>9</sup> |
| Pseudomonadota | *Enterobacter cloacae* | – | 146 ± 53 | 69 ± 21 | 5.0 × 10<sup>10</sup> ± 3.9 × 10<sup>10</sup> |
| Verrucomicrobiota | *Akkermansia muciniphila* | – | 178 ± 112 | 78 ± 24 | 8.7 × 10<sup>10</sup> ± 3.2 × 10<sup>9</sup> |
| Fusobacteria | *Fusobacterium nucleatum sp. animalis* | – | 150 ± 64 | 85 ± 24 | 1.9 × 10<sup>10</sup> ± 1.5 × 10<sup>10</sup> |
| BHI control | | | 138 ± 63 | NA | 1.4 × 10<sup>9</sup> ± 1.8 × 10<sup>9</sup> |
| MRS control | | | 130 ± 61 | NA | 6.0 × 10<sup>8</sup> ± 9.2 × 10<sup>7</sup> |

## Results

### Morphological characteristics of bEVs

Extracellular vesicles were detected in samples of 12 bacterial species by both nanoparticle tracking analysis (NTA) and transmission electron microscopy (TEM). The NTA-based mean bEV diameters ranged from 146–178 nm in gram-positive bacteria and 104–247 nm in gram-negative bacteria (Table 2). The bEV diameters were typically smaller by TEM in comparison to NTA measurements.

Based on TEM, most bEVs were likely composed of a single membrane (Fig. 1). A lipid bilayer approximately 8–10 nm thick was visible in most gram-negative bEVs, as well as in *Limosilactobacillus reuteri* and *Turicibacter sanguinis*. In the other samples, the bilayer structure was less distinct. A vesicle with double bilayer membranes was observed in *Enterobacter cloacae*. *Veillonella magna* displayed two types of bEVs: large, crumpled vesicles and small, rounded vesicles. In *Streptococcus salivarius*, most objects appeared as chain-like structures with a diameter of 20–25 nm (see top right of the image), compatible with the chains of very small vesicles previously reported in *S. pneumoniae* (60). Electron-dense material was observed in *Bacteroides fragilis, Bifidobacterium longum* sp. *suis, Enterococcus faecalis,* and *Akkermansia muciniphila* bEVs, and as central foci inside *L. reuteri* vesicles.

**Figure 1.**
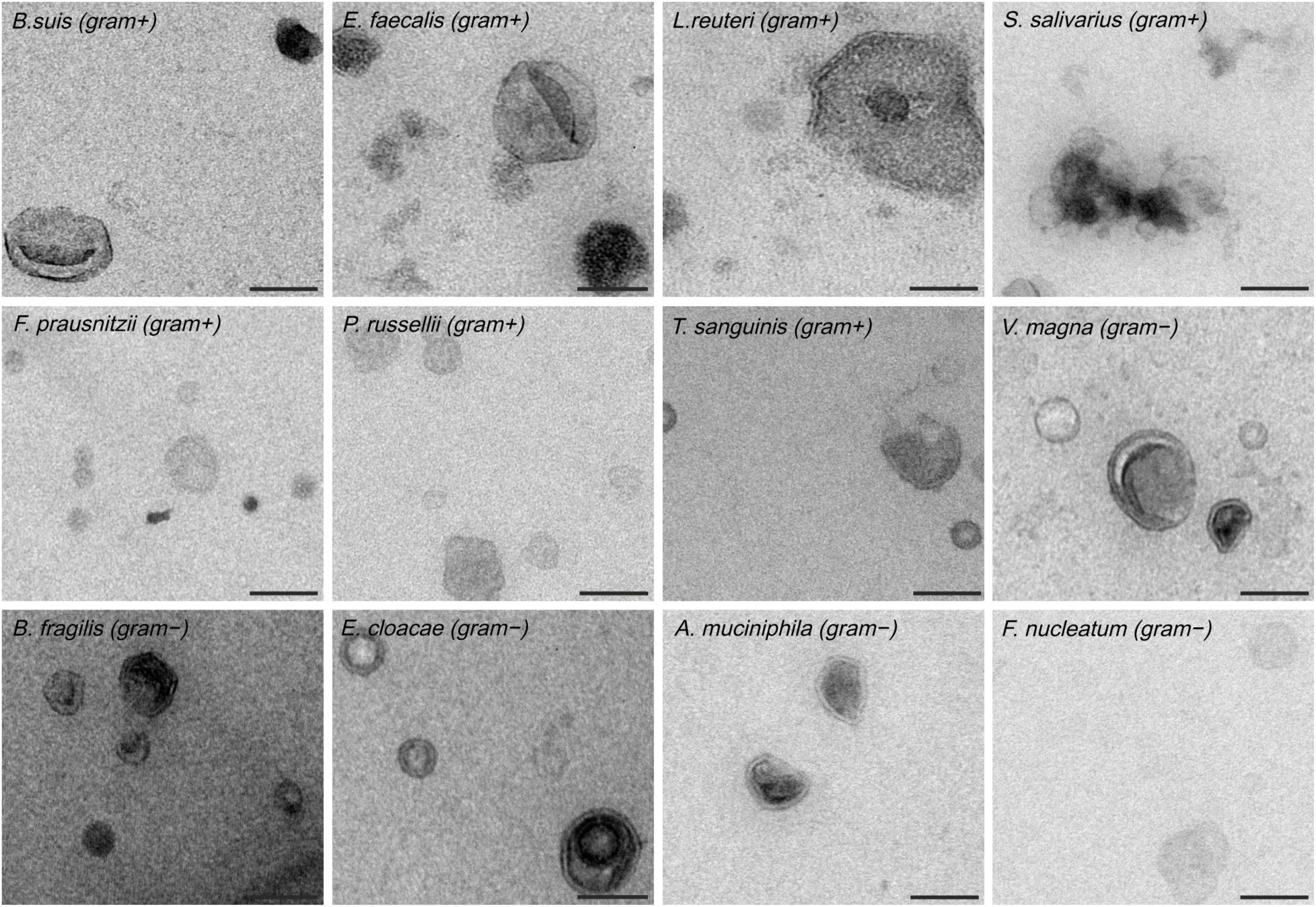
Transmission electron microscopy of extracellular vesicles from gram-positive (+) and gram-negative bacteria (−). Scale bars: 100 nm.

In the other eight species, bEVs could not be unequivocally documented, and they were excluded from further analyses. NTA-based bEV concentrations were insufficient for TEM in *E. coli*, *Flavonifactor plautii*, *Floccifex porci*, *Lactobacillus amylovorus,* and *Sutterella parvirubra*. In *Holdemanella biformis* samples, the bEV concentration was not above the level of the negative controls. TEM did not reveal EV-like particles in *Collinsella aerofaciens* and *Clostridium leptum*. *C. aerofaciens* exhibited likely protein aggregates without obvious vesicle structure. In *C. leptum* samples, we only observed long fibers with diameters ranging from 2.5–3 nm in the TEM.

### Protein content of bEVs and association of protein diversity with vesicle size

Using high-resolution LC-MS/MS, we identified 17–1297 bacterial proteins per species. These comprised 43–92% of the total MS/MS signal intensity; the remainder were animal proteins most likely derived from the culture media (Table 3). The numbers of identified bacterial proteins were associated with the NTA-measured EV concentrations (Spearman ρ = 0.59, p = 0.049). In samples with EV concentrations above 2 × 10^10^ particles/ml (Table 2), the identified bacterial proteins covered 15–49% of the reference proteomes of each species and constituted >85% of all identified proteins and >50% of the total MS/MS intensity (Table 3). In samples with lower EV concentrations, the protein coverages were lower; however, even for *F. nucleatum*, we were able to detect low-abundance proteins down to 0.3% of the total bacterial protein intensity.

**Table 3.** Number of identified bacterial proteins.

| Species | Number of bacterial proteins identified | % of the total number of proteins in samples | % of reference proteome | % of total MS/MS protein intensity |
| --- | --- | --- | --- | --- |
| *B. longum* | 63 | 50 | 3 | 9 |
| *E. faecalis* | 1269 | 92 | 46 | 94 |
| *L. reuteri* | 580 | 87 | 28 | 51 |
| *S. salivarius* | 80 | 46 | 4 | 15 |
| *F. prausnitzii* | 1297 | 94 | 49 | 91 |
| *P. russellii* | 526 | 90 | 31 | 50 |
| *T. sanguinis* | 30 | 43 | 1 | 65 |
| *V. magna* | 388 | 90 | 20 | 84 |
| *B. fragilis* | 324 | 68 | 8 | 29 |
| *E. cloacae* | 779 | 88 | 15 | 86 |
| *A. muciniphila* | 512 | 86 | 24 | 76 |
| *F. nucleatum* | 17 | 63 | 1 | 29 |

The diversity of bacterial proteins (as measured by the numbers of abundant protein orthogroups) was significantly associated with NTA-measured EV diameters (Spearman ρ = 0.67; p = 0.025).

### Predicted subcellular localizations of bEV proteins

The bacterial proteins identified in gram-positive bEV samples were predicted to be localized in the cytoplasmic membrane, cytoplasm, extracellular space, and cell wall (in the order of median relative abundances; Fig. 2). Gram-negative bEV samples contained proteins from the outer membrane, cytoplasm, extracellular space, periplasm and cytoplasmic membrane. In *F. nucleatum* bEVs, periplasmic and extracellular proteins were not observed, possibly due to the low number of identified proteins.

**Figure 2.**
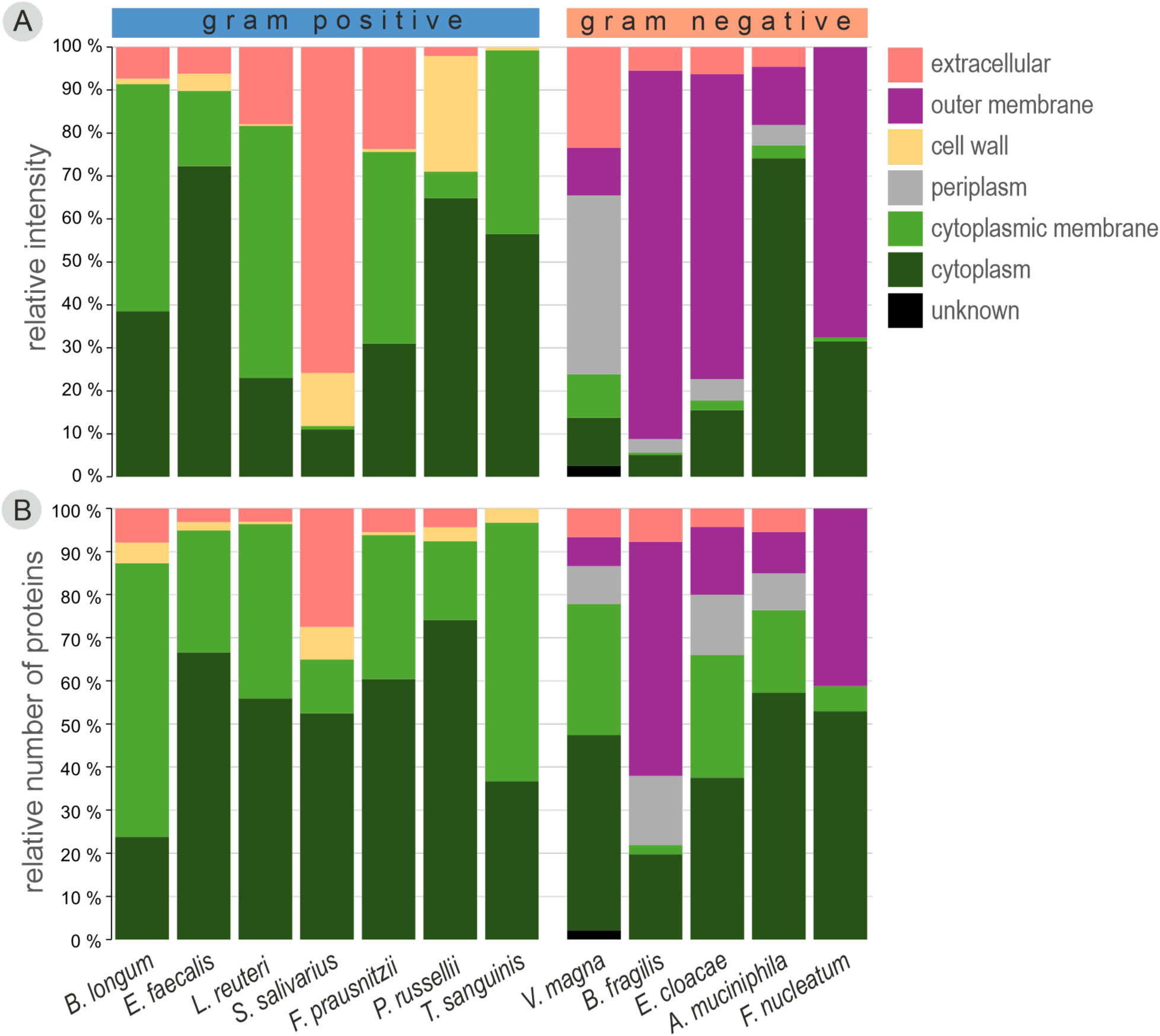
Predicted subcellular localizations of bacterial extracellular vesicle proteins. (A) relative MS/MS intensities; (B) relative numbers of identified proteins.

*A. V. magna* bEVs contained a larger share of cytoplasmic membrane and periplasmic proteins than the other gram-negative species, in terms of MS/MS signal intensity (Fig. 2A). Cytoplasmic proteins were relatively abundant in *A. muciniphila* and *F. nucleatum* bEVs. *S. salivarius* and *Peptostreptococcus russellii* bEVs contained a lower share of cytoplasmic membrane proteins compared to other gram-positive species. In *S. salivarius*, >60% of the total intensity consisted of four very abundant glucan-modifying enzymes predicted to be extracellular.

The shares of cytoplasmic membrane, cytoplasmic, and periplasmic proteins were generally higher in all species based on the number of identified proteins than by protein intensity (Fig. 2). This indicates a higher diversity of proteins in these compartments in comparison to the extracellular space, cell wall and outer membrane.

### Predicted functions of bEV proteins

The most prevalent protein functional categories included cell wall/membrane/envelope biogenesis; nutrient transport and metabolism (especially carbohydrates, amino acids, and lipids); energy production and conversion; and translation, ribosomal structure, and biogenesis (Fig. 3). Proteins for RNA and DNA processing were observed in most species. There were no major differences in the predicted protein functional categories of gram-positive and gram-negative bEVs.

**Figure 3.**
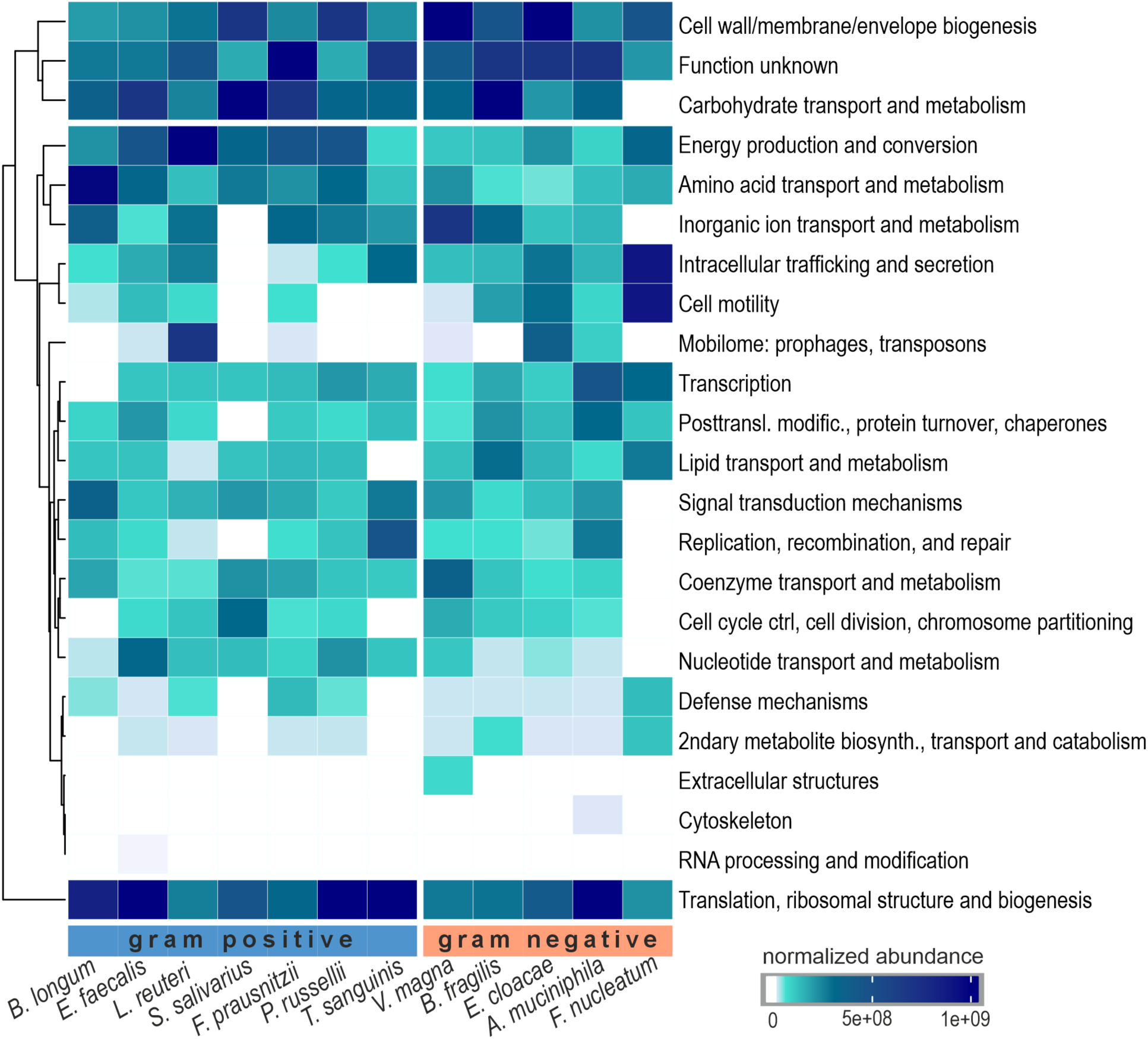
Predicted functions of bEV proteins. The heatmap shows the quantile-normalized total MS/MS intensities of clusters of orthologous groups (COG) functional categories.

Phage proteins were detected in bEVs of several species (Fig. 3). In *E. cloacae*, we observed a high abundance of P2-type temperate phage proteins, with a major capsid protein among the top 10 most abundant bEV proteins. Also in *L. reuteri*, a major capsid protein was among the most abundant proteins (>10% of the total bacterial protein intensity).

Many of the bEV proteins could not be assigned to a functional category. Among these, the *F. prausnitzii* protein WP_015537825.1 (microbial anti-inflammatory molecule, MAM) and the *A. muciniphila* proteins WP_197738471.1 (Amuc_1100), and WP_012420447.1 (Amuc_1409) are known to mediate host-microbe interactions (61).

### Correlations between bEV proteome compositions, phylogeny and gram type

The bEV proteome compositions primarily reflected phylogenetic relationships between the bacterial species (Fig. 4). The *V. magna* proteome grouped with the other Bacillota (Firmicutes) despite the difference in gram type. The cophenetic correlation coefficient between the proteomic and phylogenetic trees was 0.46, indicating a moderate positive correlation between bEV protein composition and bacterial phylogeny. The Fowlkes-Mallows index showed significant non-random similarity between two trees in four out of nine numbers of clusters (two to five clusters); the similarity was random only in one case with nine clusters. Most of the protein orthogroups detected in more than one species (60% of all orthogroups) were shared between gram-positive and gram-negative species. However, *B. longum* of the Actinomycetota grouped with Bacillota, possibly indicating effects of gram type.

**Figure 4.**
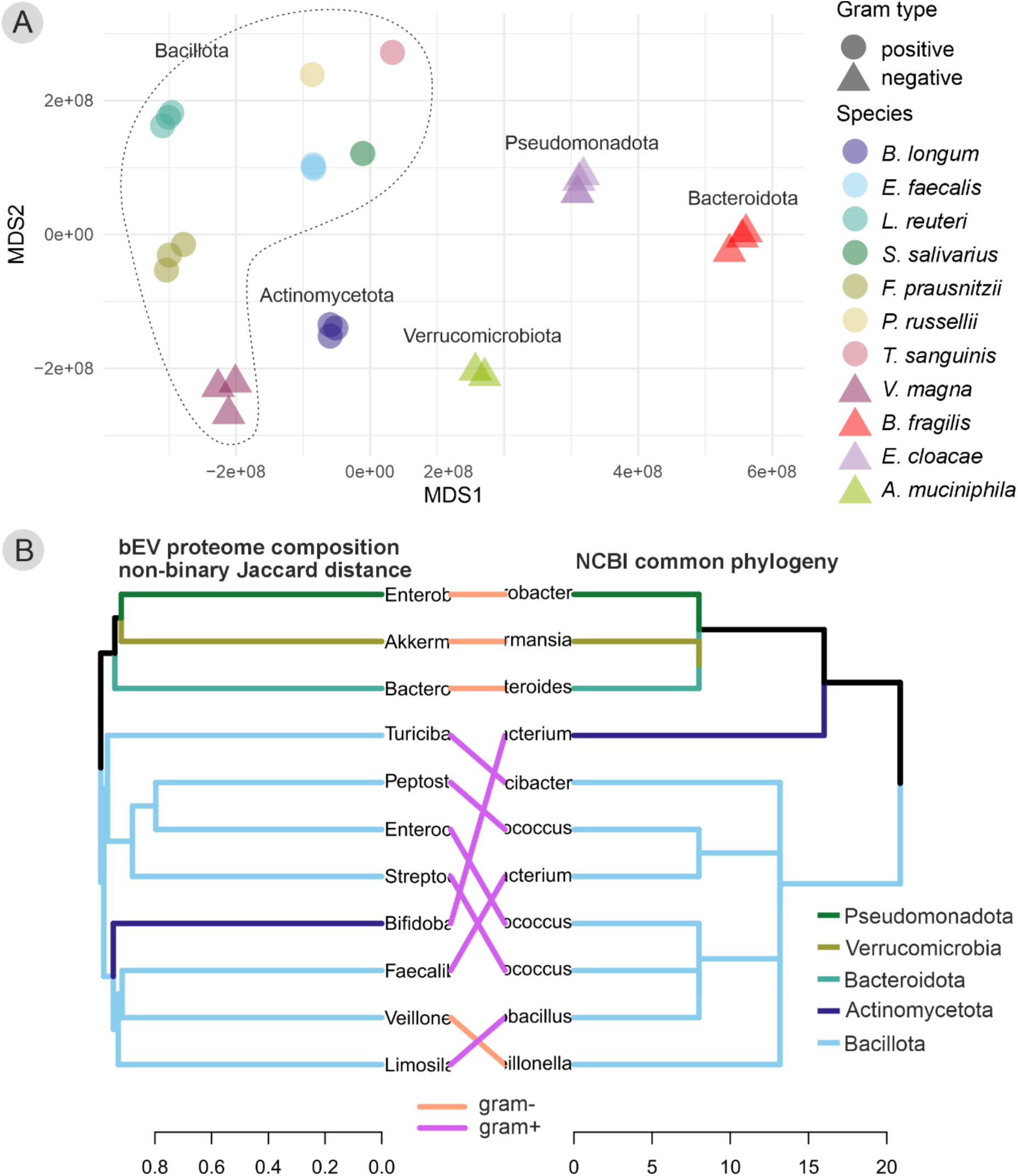
bEV proteome compositions and phylogenetic relationships. (A) nMDS ordination using Euclidean distance matrix of proteome composition (Stress = 0.1). Replicate samples are shown. For some of the species, replicates were pooled to increase LC-MS/MS intensity. The dotted line shows the species of the Bacillota (Firmicutes) phylum. (B) Comparison of proteome composition tree based on bEV protein similarity (complete linkage, non-binary Jaccard distances) and the NCBI common phylogeny tree, which reflects the expected evolutionary relationships derived from nucleotide sequence data. Tree branch colors indicate phyla; connector line colors indicate gram type.

The *F. nucleatum* proteome was not included in these phylogenetic analyses due to the small number of identified proteins.

### Conserved and abundant bEV proteins

Conserved protein orthogroups were defined as those present in at least five gram-positive species and in four gram-negative species. We identified 14 orthogroups that were conserved in bEVs from both gram-positive and gram-negative bacteria (Fig. 5A). These were predominantly cytoplasmic proteins (Fig. 5B). Glyceraldehyde-3-phosphate dehydrogenase (GAPDH), elongation factor Tu (Ef-Tu), DNA-directed RNA polymerase subunit beta, and several ribosomal proteins had the highest median MS/MS signal intensities. Although these proteins were abundant in some species, the medians were all below 2% of total MS/MS intensity.

**Figure 5.**
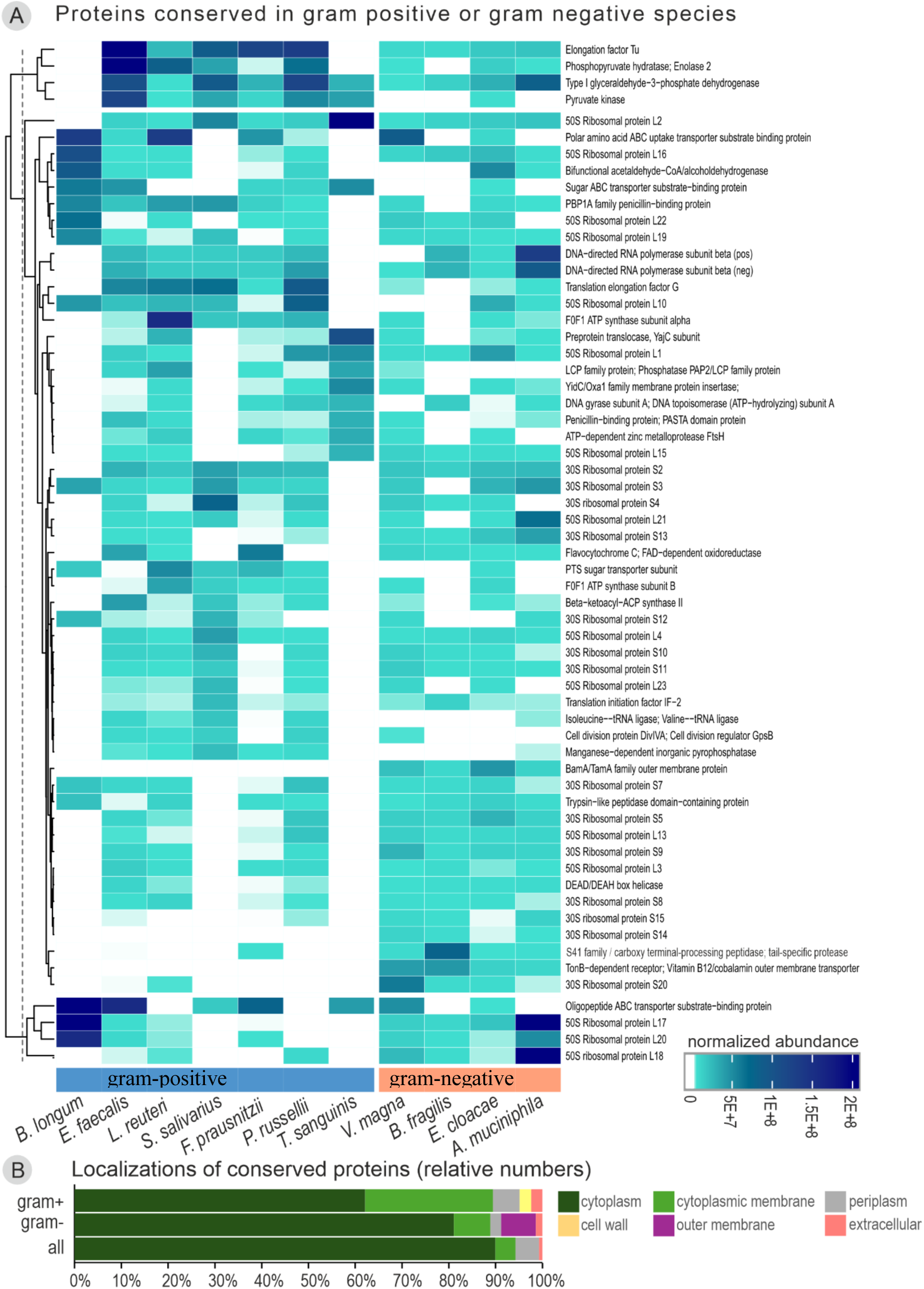
Conserved bEV proteins. A. A heatmap of normalized MS/MS intensities of bEV proteins conserved in gram-positive or gram-negative bacteria. B. The predicted subcellular localizations for bEV proteins conserved in gram-positive bacteria, gram-negative bacteria, and across all species (as relative numbers of identified proteins).

An additional 29 orthogroups were conserved in gram-positive bEVs only (Fig. 5A). Among all the proteins conserved in gram-positive bEVs, the most abundant were GAPDH, Ef-Tu and PBP1A family penicillin-binding proteins (PBPs).

Eighteen orthogroups were conserved in gram-negative bEVs only. The most abundant among all the proteins conserved in gram-negative bEVs included TonB-dependent receptor, S41 family peptidase, GAPDH, DNA-directed RNA polymerase, and some ribosomal proteins. TonB-dependent receptor, BamA/TamA family outer membrane protein assembly factors, and the 30S ribosomal protein S14 were found exclusively in gram-negative bEVs.

We assessed the ten most abundant proteins in each species. A majority of them were species-specific, with only a small proportion shared across several species (Fig. 6A). The top 10 proteins accounted for 21–94% of the total MS/MS signal intensity per species and were often predicted to be involved in cell envelope biogenesis, carbohydrate processing or translation; however, in many cases, the functions of these proteins are unknown.

**Figure 6.**
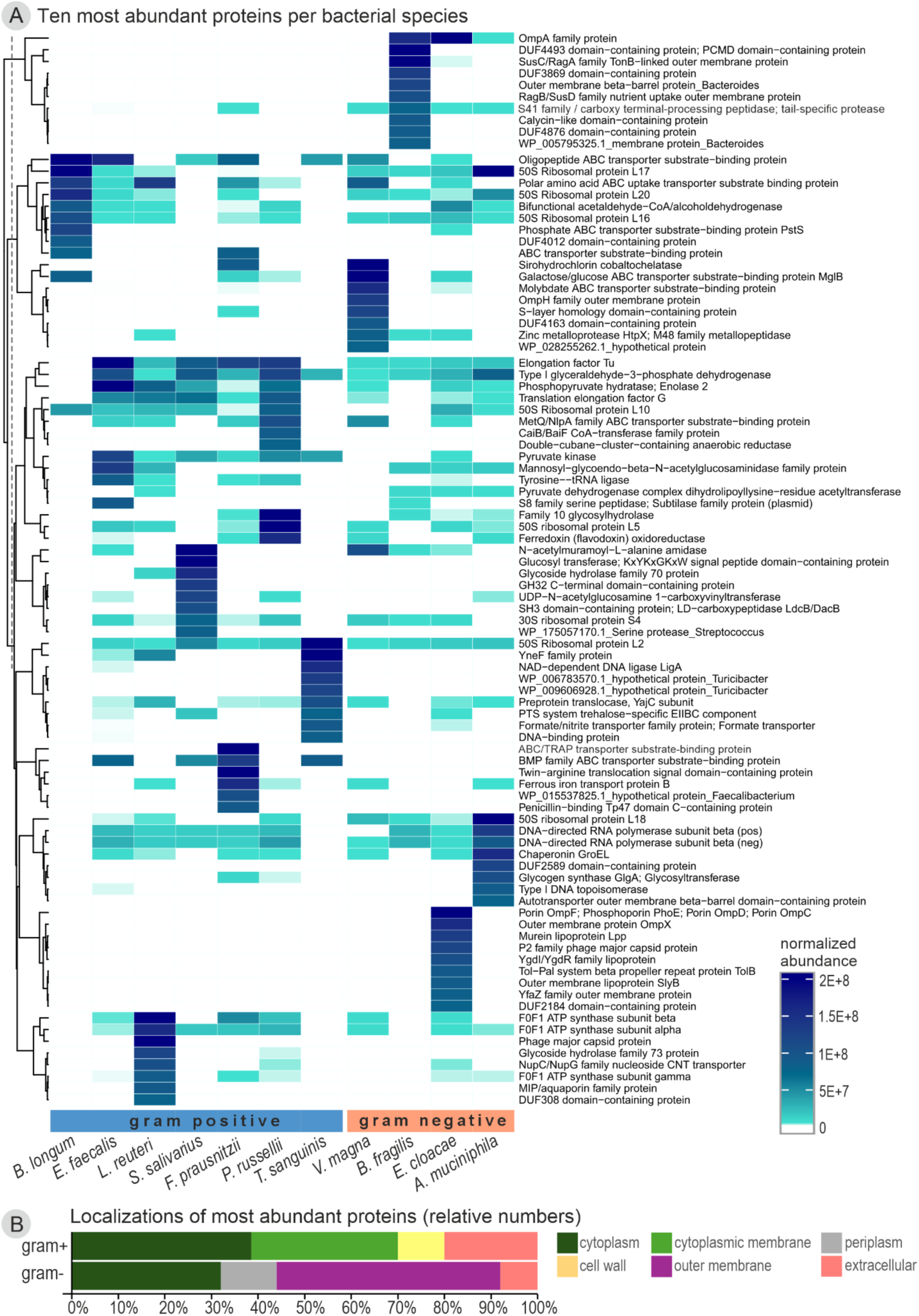
Most abundant bEV proteins. A. Quantile-normalized abundances of proteins which were among the 10 most abundant in at least one bacterial species. B. Predicted subcellular localizations for the most abundant bEV proteins in gram-positive and gram-negative bacteria (as relative numbers of identified proteins).

In *V. magna*, the most abundant protein was a cobalt chelatase involved in vitamin B_12_ production (Fig. 6A). In *L. reuteri*, the abundant proteins were mostly related to ATP synthesis.

*A. muciniphila* bEVs were abundant in chaperonin GroEL, while in *B. longum*, the PBP1A protein dppA accounted for over 23% of total bacterial protein intensity. *T. sanguinis* bEVs were abundant in DNA repair proteins and *P. russellii* bEVs in oxidative stress proteins. Phage components were among the ten most abundant bEV proteins in *L. reuteri* and *E. cloacae*. In *T. sanguinis*, >20% of total bacterial protein intensity consisted of a single cytoplasmic membrane protein with unknown function.

In gram-positive bEVs, the majority of the most abundant proteins were predicted to be localized in the cytoplasm, cytoplasmic membrane, or the extracellular space (Fig. 6B). In gram-negatives, the most abundant proteins were in outer membrane and cytoplasm, followed by periplasm and the extracellular space (Fig. 6B).

### Comparison to previously published bEV data

We compared our data to previously published *in vitro* bEV proteomics data (available for five species) and fecal EV metaproteomics data (two datasets in which sufficient numbers of peptides could be assigned at the genus level). The bacterial strains and culture media used in the *in vitro* studies differed, except for *E. cloacae*, which was the same strain.

In *B. longum,* the subcellular distribution of bEV proteins was similar, despite the difference in the number of identified proteins (63 in our data versus 552 in the previous study) (Fig. 7A) (31). The intensity shares of the major protein functional categories (>5%) were also similar, except for defense mechanisms and the posttranslational modification functional category, which were much less abundant in our data (Fig. 7B).

**Figure 7.**
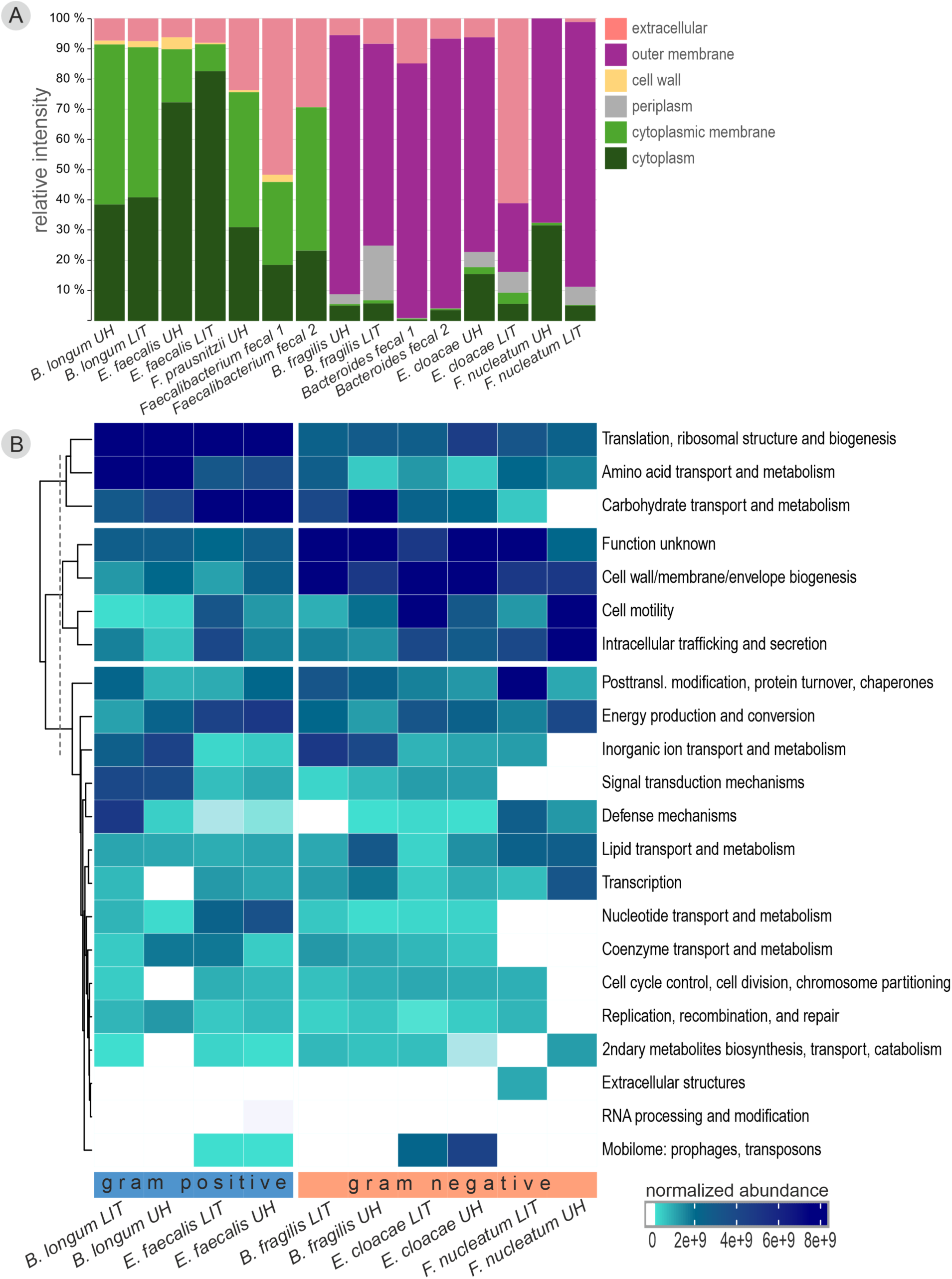
Comparison of predicted subcellular localizations. (A) and functions (B) of bEV proteins in our a data (UH) and previously published data (LIT; literature). *B. longum* (*31*)*, E. faecalis* (62), the *Faecalibacterium* genus in fecal metaproteomics data 1 & 2 (57, 58)*, B. fragilis* (27), the *Bacteroides* genus in fecal metaproteomics data 1 & 2 (57, 58), *F. nucleatum* (*64*)*, E. cloacae* (63).

There was also a high degree of similarity in protein subcellular localizations and functional categories in the *E. faecalis* protein datasets (Fig. 7A) (62). With >1100 identified proteins in both datasets, all functional categories with an intensity above 5% matched between the two datasets. The differences in the categories of posttranslational modifications and defense mechanisms were mainly the result of a few individual proteins having higher intensity in either dataset, while the number of proteins per category was similar.

For *B. fragilis,* the protein subcellular localizations were relatively similar, with a higher proportion of proteins in the outer membrane and fewer in the periplasmic space in our data (Fig. 7A) (27). There was also a good match of functional annotations, with some exceptions (Fig. 7B). Carbohydrate transport and metabolism had a lower total number of proteins in our dataset but contained one protein with very high relative intensity (34%) that was not present in the previously published dataset. In the category of amino acid transport, the previously published dataset had only a slightly higher number of proteins, but several of them had higher individual intensities than the sum intensity of all these proteins in our dataset.

*E. cloacae* datasets showed the largest variation in protein localizations between our data and previously published data (Fig. 7A) (63). The share of outer membrane proteins was much higher in our data, and extracellular proteins were lower. The functional annotations were similar with two major exceptions (Fig. 7B): cell motility proteins were much more abundant in the previously published dataset, due to the very high intensity (50%) of the flagellin FliC. The cell wall/membrane/envelope biogenesis category was much more abundant in our data, with several outer membrane proteins (Omp) comprising >40% of total intensity.

For *F. nucleatum*, we detected a smaller number of proteins than in the previously published dataset (17 versus 98) (64). The share of cytoplasmic proteins was higher and outer membrane proteins lower in our data (Fig. 7A). The relative intensities of several of the functional categories also differed, albeit several with lower intensity still had similar levels in both datasets (Fig. 7B). Major differences in the categories cell motility-intracellular trafficking and posttranslational modification, both consisting of only a few proteins in each dataset, were due to vastly higher intensities of one or two proteins.

In the fecal EV metaproteomics data, the localizations of proteins assigned to the genus *Faecalibacterium* (57, 58) were quite similar to our *F. prausnitzii* data, although the fecal data had a higher share of extracellular proteins (Fig. 7A). The fecal *Bacteroides* proteomes (57, 58) contained fewer cytoplasmic and periplasmic proteins than both *in vitro* datasets; this may be in part because conserved proteins often cannot be assigned to a specific genus and were therefore omitted in our analysis. We did not compare the functional annotations to the fecal EV metaproteomics datasets, as species-level annotation was not possible.

## Discussion

We characterized the extracellular vesicle proteomes of 12 commensal bacterial species, representing the six major phyla of the mammalian intestinal microbiota. Predicted protein functions suggest possible roles for commensal bEVs in nutrient processing and host immune system modulation. We identified conserved proteins that may be used as markers for differentiating bacterial versus host EVs. Of note, this is the first report of bEVs secreted by *V. magna*, representing an exceptional class of gram-negative Bacillota (Firmicutes), as well as for *T. sanguinis* and *P. russellii*. We provide the first publicly available proteome of *F. prausnitzii* EVs.

The structures and proteomes of bEVs from gram-positive and gram-negative bacteria reflected the distinct architectures of their parent cells (1). In gram-positive bacteria, a thick peptidoglycan cell wall surrounds the cytoplasmic membrane. They secrete cytoplasmic membrane vesicles (CMVs) by membrane blebbing or explosive cytoplasmic membrane vesicles (ECMVs) by bubbling cell death, through holes degraded in the cell wall (1). In our data, gram-positive bEVs were often irregularly shaped and contained proteins primarily localized to the cytoplasmic membrane and cytoplasm, with varying amounts of extracellular and cell wall proteins.

Gram-negative bacteria have a cytoplasmic membrane and an outer membrane separated by a periplasmic space containing a thin peptidoglycan layer. Here, bEV biogenesis by blebbing produces two types of vesicles: outer membrane vesicles (OMVs), which have a single membrane and periplasmic cargo, but no cytoplasmic cargo; and outer-inner membrane vesicles (OIMVs) with a double membrane and cytoplasmic material. Biogenesis by explosive cell lysis generates explosive OMVs (EOMVs) and explosive OIMVs (EOIMVs), both containing cytoplasmic material, with EOIMVs having more than one vesicle in a larger vesicle (1). In our data, the gram-negative bEVs were mostly spherical with a distinct bilayer membrane, and contained outer membrane, cytoplasmic, extracellular, periplasmic and cytoplasmic membrane proteins. The high share of cytoplasmic proteins suggests biogenesis by explosive cell lysis; thus, the gram-negative bEVs were likely a mixture of EOMVs and OMVs in most cases. *A. muciniphila* samples had a very high abundance of cytoplasmic proteins, in agreement with a previous report (21).

*V. magna* may produce EVs from both the outer membrane and the cytoplasmic membrane: electron micrographs showed two different types of single-membrane vesicles, and the samples contained equal shares of outer membrane and cytoplasmic membrane proteins. This is unexpected, as gram-negative bacteria are not known to secrete vesicles composed solely of cytoplasmic membrane (1). This warrants further research on EV biogenesis mechanisms in Negativicutes. In *E. cloacae* bEVs, a rare double membrane vesicle was observed, consistent with a previous report (63). Electron-dense material was observed in *B. fragilis* and *A. muciniphila* bEVs, and as central foci in *L. reuteri* bEVs, possibly representing cytoplasmic components (65).

bEV diameters did not show marked differences between gram types or taxa. They were largely consistent with previous studies, including *B. fragilis* (18), *E. cloacae* (63), *S. salivarius* (66), *B. longum* (5)*, E. faecalis* (67), *L. reuteri* (68) and *A. muciniphila* (69). Our *F. prausnitzii* bEVs were larger than those reported by Ye et al. (70), which may be due to technical factors such as isolation and purification methods or potential strain differences (71). Electron microscopy typically showed smaller bEV sizes compared to NTA. This may be because NTA measures the hydrodynamic diameters rather than exact membrane-bound dimensions, the lower sensitivity of NTA to small particles, or the shrinkage during the EM fixation (72, 73).

Our proteomics data likely covers most of the major bEV proteins, even in the species with the lowest vesicle yields. A 100-nm bEV is expected to contain approximately 4000 protein molecules, based on consensus estimates of cytoplasmic and membrane protein concentrations (74–76). In all proteomes with >60 identified proteins, the intensity ratio of the most abundant vs. least abundant proteins was above 4000, implying that theoretically, we detected down to one molecule per a vesicle. The bEV proteomes with >500 identified proteins covered 15–50% of the reference genomes. However, this represents the pooled proteome of a large number of vesicles, and a single vesicle would probably contain a smaller number of detected proteins. If only proteins with relative intensity above 1/4000 are considered (theoretically representing the protein packaging capacity of a single 100-nm vesicle), the genome coverage was 4–16%. As expected, bEVs with larger diameters were associated with higher diversities of abundant proteins.

Functional annotation of bEV proteomes revealed abundant cell envelope-modifying proteins, suggesting involvement in vesicle biogenesis by blebbing. Peptidoglycan biosynthesis and remodeling proteins were present in both gram-positive and gram-negative bEVs, and included Mur ligases and transferases, penicillin-binding proteins (PBPs), and MreB-C rod-shape determining proteins (77, 78). These have been previously reported for bEVs of *E. cloacae* (79) and several other species (80). Bacterial growth and division involve the synthesis of new peptidoglycan and the degradation of old peptidoglycan by hydrolases (78, 81). This creates weakened regions that permit the membrane to bleb from the cell (2, 82). The enrichment of the peptidoglycan biosynthesis and remodeling proteins at sites of active cell wall turnover promotes their incorporation into bEVs (78).

Phage-mediated bEV biogenesis has been demonstrated in gram-positive and gram-negative bacteria (1, 83, 84). We identified phage proteins and the recombinase A (RecA) in bEVs from *E. cloacae, L. reuteri*, *A. muciniphila*, *F. prausnitzii*, *E. faecalis,* and *V. magna*. Major capsid proteins were abundant in *E. cloacae* and *L. reuteri* bEVs; small amounts of phage holins were detected in these species and *F. prausnitzii*. Genotoxic stress can activate RecA, which induces the expression of lytic genes such as endolysins and holins from prophages. Holins create holes in the cytoplasmic membrane, allowing endolysins to access and weaken the peptidoglycan layer. This promotes vesicle biogenesis by bubbling cell death in gram-positive bacteria or cell lysis in gram-negative bacteria (83–87). Phage proteins were reported in *E. cloacae* and *E. faecalis* bEVs previously (63, 79).

Nutrient metabolism and transport proteins were the most common functional group in almost all species, and were often highly abundant. Cobalt chelatase, which is involved in vitamin B_12_ metabolism, was abundant in bEVs of *V. magna* and *F. prausnitzii*. These enzymes provide vitamin B_12_ to other microbes and the host (88). GAPDH, the key glycolytic enzyme with additional roles in iron uptake and membrane trafficking (75, 76, 101), was present in bEVs from most species. Glycoside hydrolases were abundant in *S. salivarius* bEVs and present in lower amounts in other species. They break down complex polysaccharides, providing nutrients for both the producing bacteria and other microbes (11, 89, 90). ABC transporters responsible for the transport of iron, oligopeptides, and amino acids across membranes (10, 91) were abundant in *B. longum* and widely detected across species. TonB-dependent receptors, which facilitate the uptake and delivery of nutrients (10, 92), were found in the bEVs of all gram-negative species. SusC/RagA and SusD/RagB family nutrient uptake outer membrane proteins, involved in the uptake of complex polysaccharides (93), were exclusively detected in *B. fragilis* bEVs.

The bEV proteomes of all species included known immunomodulators (61). The elongation factor EF-Tu has cell surface moonlighting functions in adhesion and immune system modulation (94). It was detected in bEVs from nine species and was particularly abundant in *E. faecalis* and *P. russellii*. The OmpA outer membrane porin family of commensal bacteria can promote intestinal homeostasis by inducing IL-22 (95). They were present in all gram-negative bEV samples, with high abundances in *E. cloacae* and *B. fragilis*. The *F. prausnitzii* microbial anti-inflammatory molecule (MAM) downregulates NF-κB signaling in epithelial cells (96, 97). The chaperonin/heat shock protein GroEL, known for its immunoregulatory activity (98), was especially abundant in *A. muciniphila* bEVs. The *A. muciniphila* bEVs also contained less abundant proteins that have been reported to modulate gut barrier integrity (99), epithelial stem cells (100) and GLP-1 secretion (101).

The functions of many abundant bEV proteins are unknown, especially in *T. sanguinis*, *F. prausnitzii*, and *B. fragilis*. This limits our understanding of the potential roles of bEVs and encourages experimental studies on these proteins.

The bEV proteome compositions primarily reflected phylogenetic relationships of the producing bacteria. This was especially evident in the *V. magna* bEV proteome, which clustered with other Bacillota bEVs despite the difference in gram type. Thus, bEVs largely inherit the proteome compositions of their parent cells, regardless of the differences in biogenesis mechanisms. This suggests the majority of proteins were loaded into bEVs non-selectively.

To identify potential markers differentiating bEVs from host EVs, we assessed for conserved bEV proteins. Although the majority of the abundant bEV proteins were species-specific, we detected 14 orthologous proteins in the bEVs of most gram-positive and gram-negative bacteria. These were primarily predicted to be cytoplasmic, and as such, not ideal as universal bEV markers, since OMVs of gram-negative bacteria lack cytoplasmic proteins (1). The most abundant of the broadly conserved bEV proteins included Ef-Tu, GAPDH, DNA-directed RNA polymerase subunit beta, and certain ribosomal proteins. EF-Tu is listed among the top 50 most conserved bEV proteins deposited in the EVpedia database (102) and may be partially localized on the cell surface (94). GAPDH has been proposed as a target for vaccines against bacterial infections due to its abundance, partial membrane localization, involvement in virulence, and limited homology with host GAPDH (103). The 30S ribosomal protein S4 was found in most bacterial species and does not have mitochondrial orthologues present in animal EVs.

TonB-dependent receptor was the most prevalent outer-membrane protein and is a potential marker for gram-negative bEVs. BamA/TamA family outer membrane protein and 30S ribosomal protein S14 were also exclusively detected in gram-negative bEVs, although less abundant. PBP1A is a potential marker for gram-positive bEVs, although related proteins were also present in some gram-negative bEVs.

bEV proteomes are likely affected by experimental methods, bacterial growth conditions, and the strain of the producing bacteria (1). However, the predicted localization patterns of proteins observed in our study were consistent with those reported in the literature, including *in vitro* bEVs from *F. nucleatum* (64), *B. fragilis* (27), and *E. faecalis* (62), as well as *Bacteroides* and *Faecalibacterium* proteomes extracted from fecal EV metaproteomes (57, 58). This suggests that the types of bEVs obtained from different experimental procedures, growth conditions, and bacterial strains are reproducible. The lower proportion of cytoplasmic proteins detected in the fecal bEV data may be attributed to their conserved nature, which prevents the assignment of peptides to specific taxa.

There was also a general similarity in the predicted functions between our proteomic data and previously published *in vitro* data. This was particularly evident in *E. faecalis* bEVs, with both studies detecting large numbers of proteins (62). In other species, observed differences were mostly due to individual proteins present at highly different intensities. For example, our *B. fragilis* data included a highly abundant outer membrane protein with unknown function that was absent in the previous study (27).

In some species, we detected fewer proteins compared to the literature datasets, likely due to low bEV concentrations. *B. longum* bEVs contained considerably fewer proteins in our study (31). This probably affected the detection of proteins of the smaller functional categories, although the intensity shares of the subcellular localizations and the major functional categories were similar. For *F. nucleatum* bEVs, despite differences in culture media, strain, and the very low number of identified proteins in our dataset, there were similarities in both cellular localization patterns and the distribution of protein functional categories (64).

### Limitations of the work

The culture conditions were not optimized for each species, as we used a standardized protocol, and the growth phases of the cultures were not controlled. These may affect the yields, types, and compositions of bEVs (104). The observed bEV concentrations probably do not indicate the EV production capacity of these species *in vivo*. In bEV extraction, we used a 0.22 µm filter, which may have excluded some of the larger bEVs, but it was necessary to reliably remove bacterial cells. Protein amounts injected in LC-MS/MS were not equalized, as we aimed for maximal resolution for each species. The number of identified bacterial proteins is therefore not directly comparable between species, but the relative ratios of the subcellular localizations and main functional categories can be compared.

## Conclusions

Our study provides an overview of commensal bEV proteomes across the major phyla of the mammalian intestinal microbiota.

The bEV compositions reflected the envelope structures of gram-positive and gram-negative bacteria. The gram-negative Bacillota *V. magna* (and possibly other Negativicutes) may have an exceptional ability to produce both outer membrane and cytoplasmic membrane vesicles.

Peptidoglycan modifying enzymes observed in several species and bacteriophage proteins very abundant in *E. cloacae* and *L. reuteri,* probably mediate bEV biogenesis.

Commensal bEVs are likely involved in nutrient metabolism and transport as well as in the modulation of the host immune system. The functions of many abundant bEV proteins are unknown.

Elongation factor Tu and GAPDH are potential generic markers for bEVs. TonB-dependent receptors and PBP1A may be used as markers for gram-negative and gram-positive bEVs, respectively.

Our comparison to previous bEV studies suggests that bEV structures and their proteomic profiles are largely reproducible, despite differences in bacterial growth conditions, bEV isolation protocols, and analysis methods.

## Author contributions

**RNM:** data analysis (lead), visualization (supporting), writing (lead). **AM:** proteomics and NTA data analysis (lead), visualization (lead), writing (supporting). **TPM:** data analysis and writing (supporting). **MK:** TEM and NTA data analysis and writing (supporting). **MS:** proteomics laboratory analysis (lead), writing (supporting). **MH:** bacterial strains (supporting), writing (supporting). **AK:** metaproteomics data analysis (supporting), writing (supporting). **SM:** methodology and writing (supporting). **JR:** resources, conceptualization and writing (supporting). **TRL:** resources and writing (supporting). **TAN:** proteomics laboratory analysis (lead), proteomics data analysis and writing (supporting). **MN:** conceptualization, funding acquisition, methodology, resources, supervision, writing (lead), data analysis, visualization (supporting).

## Acknowledgments

We thank chief laboratory technician Kirsi Lahti for performing bacterial culture, and Dr. Ulla Hynönen, Dr. Silja Åvall-Jääskeläinen, Dr. Kaisa Hiippala, Dr. Reetta Satokari and prof. Willem de Vos for expert consultation. We acknowledge the services of the University of Helsinki: the EV Core Facility in Viikki for EV isolation and NTA analysis, the HiPREP Core in the FIMM Technology Centre supported by HiLIFE and Biocenter Finland for electron microscopy, the Electron Microscopy Unit of the Institute of Biotechnology for providing the facilities, the Clinical Microbiology Laboratory of the Veterinary Teaching Hospital (YESLAB) for the MALDI identifications of the *in vitro* cultured bacteria, and the DNA Sequencing and Genomics Laboratory (supported by HiLIFE and Biocenter Finland funding), Institute of Biotechnology for DNA sequencing.

Mass spectrometry-based proteomic analyses were performed by the Proteomics Core Facility, Department of Immunology, University of Oslo/Oslo University Hospital, which is supported by the Core Facilities program of the South-Eastern Norway Regional Health Authority. The core facility is a member of the National Network of Advanced Proteomics Infrastructure (NAPI), funded by the Research Council of Norway INFRASTRUKTUR-program (project number: 295910).

